# Pulmonary delivery of antigen-enhanced BCG overcomes safety barriers in immunocompromised hosts and protects against TB in the absence of adaptive immunity

**DOI:** 10.64898/2026.07.05.736631

**Authors:** Harindra Darshana Sathkumara, Guangzu Zhao, Andrew Calcino, Munish Puri, Socorro Miranda-Hernandez, Yide Wong, Julia Seifert, Matt A. Field, Roland Brosch, Andreas Kupz

## Abstract

The low efficacy of Bacille Calmette-Guérin (BCG) in preventing pulmonary tuberculosis (TB) underscores the need for improved TB vaccines. Recombinant BCG (rBCG) strains secreting the virulence-associated effector molecule ESAT-6 from *Mycobacterium tuberculosis* (*Mtb*) markedly improve efficacy and immunogenicity in animal models of TB but have been considered unsuitable for clinical translation due to safety concerns identified in intravenous SCID mouse models. Here, we demonstrate that pulmonary delivery fundamentally reshapes the safety and protective efficacy of the ESAT-6-secreting rBCG strains, BCG::RD1 and BCG::ESAT6-PE25SS. In sharp contrast to intravenous delivery, pulmonary administration was markedly better tolerated, improved survival, and reduced systemic dissemination and brain pathology of severely immunocompromised mice. Strikingly, pulmonary rBCG vaccination also conferred superior protection against aerosol *Mtb* challenge in wild-type, type 2 diabetic, and adaptive immunity-deficient *Rag1^⁻/⁻^* and *Rag2^⁻/⁻^Il2rg^⁻/⁻^* mice, with BCG::RD1 showing the strongest adaptive immunity-independent protection. Mechanistically, pulmonary rBCG vaccination promoted lung innate immune activation, expansion of myeloid-biased progenitors in the bone marrow, and enhanced antimycobacterial activity of macrophages, consistent with trained innate immunity. Collectively, these findings reveal that pulmonary vaccination largely overcomes safety concerns of ESAT-6-secreting rBCG strains and provide preclinical evidence for a viable strategy to improve protection against TB in immunocompromised individuals.

## Introduction

Tuberculosis (TB) remains the leading cause of infectious mortality causing over 10.8 million new infections and approximately 1.25 million deaths annually, with low- and middle-income countries disproportionately affected (1). The Bacillus Calmette-Guérin (BCG) vaccine, the only licensed TB vaccine, has been in use for over a century and remains widely administered, yet its efficacy is highly variable (2). While BCG provides protection against severe forms of TB in infants, its effectiveness in preventing pulmonary TB in adults is inconsistent and wanes over time (3). This variability in protection has been a major hurdle for global TB control, especially in endemic regions where TB transmission remains high, and no effective booster strategy has been implemented. Despite decades of effort, only a handful of TB vaccine candidates have progressed to late-stage clinical trials, and several have failed to surpass BCG in efficacy (4, 5).

One of the critical challenges in TB vaccine development is the complexity of *Mycobacterium tuberculosis* (*Mtb*) pathogenesis, which requires a coordinated interplay between adaptive and innate immune responses for effective control. Like many other bacterial pathogens, *Mtb* has evolved sophisticated mechanisms to evade host immunity and persist within macrophages, leading to chronic infection (6, 7). Live attenuated vaccines (LAVs) are therefore considered as promising TB vaccine candidates due to their ability to mimic natural infection and stimulating a broad and durable immune response (8). However, LAVs can pose safety concerns for immunocompromised individuals, as their inability to mount effective immune responses can lead to uncontrolled bacterial replication and dissemination (9). This is particularly relevant for populations such as people living with HIV with low CD4^+^ T-cell counts, individuals with congenital immunodeficiencies (e.g., SCID), and those undergoing immunosuppressive therapies. For that reason, LAV candidates are required to be assessed for safety in immunocompromised mice during pre-clinical development (10).

BCG differs from *Mtb* in multiple genomic and functional aspects, including the absence of the region of difference 1 (RD1) genomic region, which encodes an essential part of the ESX-1 type VII secretion system that secretes key virulence factors such as ESAT-6 and CFP-10, and whose absence contributes to its attenuated virulence and altered immune profile (11–13). While recombinantly reintroducing *Mtb* RD1 antigens into BCG (as seen in BCG::RD1) has experimentally enhanced immunogenicity and protection, this modification has also increased virulence, particularly in immunocompromised models, previously rendering such strains unsuitable for clinical application (14, 15). Current preclinical safety assessments of TB vaccines, however, may not fully capture real-world risks, as these models often employ very high-dose, intravenous administration of vaccines in severely immunocompromised mice (i.e., SCID, *Rag2^-/-^Il2rg^-/-^*) that do not accurately reflect human scenarios (16, 17). Moreover, this approach may also mask the beneficial aspects of some LAVs by disproportionately amplifying their persistence and pathogenicity in this highly sensitive model, thereby limiting the ability to assess their potential efficacy and safety in immunocompetent populations (18).

Moreover, beyond HIV coinfection; the prime risk factor for TB, type 2 diabetes (T2D) has emerged as a significant risk, increasing susceptibility to infection by 3-4 times due to chronic inflammation and immune dysregulation (19, 20). Interestingly, we have recently shown that, despite the previously reported RD1-associated safety concerns, pulmonary/mucosal administration of BCG::RD1 in TB/T2D comorbid mice was well tolerated and provided superior protection against aerosol *Mtb* challenge (21). These findings suggested that mucosal BCG::RD1 vaccination may play a role in shaping local immune responses in the lung, potentially inducing both adaptive and innate cell activity.

In the current study, we investigated the innate immune mechanisms that drive the superior anti-TB immunity in preclinical models of immunosuppression using BCG::RD1 and the recombinant BCG (rBCG) strain BCG::ESAT6-PE25SS (hereafter referred to as PE25SS), which selectively secretes ESAT-6 via ESX-5, a secretion pathway naturally present in BCG (22). We show that improved protection after mucosal rBCG vaccination extends to mice lacking all adaptive immunity and is driven by increased bone marrow myelopoiesis, distinct transcriptional reprograming in lung innate cells and superior innate training. These findings highlight that ESAT-6-secreting BCG strains, when delivered directly into the lung, drive potent innate immune mechanisms in the lung, compensating for the absence of adaptive immunity. To our knowledge, this study provides the first evidence that pulmonary delivery can overcome safety barriers of previously excluded “virulent” rBCG strains and unlock their protective potential through innate immune priming.

## Results

### Pulmonary delivery overcomes the safety concerns of ESAT-6-expressing rBCG in immunocompromised mice

Despite being among the most efficacious live attenuated TB vaccine candidates described to date, BCG::RD1, an RD1-complemented BCG strain, has previously been deemed unsuitable for clinical translation because of safety concerns observed in highly sensitive SCID mice (12, 14). Given our previous finding that mucosal BCG::RD1 was well tolerated in vulnerable diabetic mice (21), we hypothesized that the route of administration could be a critical determinant of safety in immunosuppressed hosts. To directly address this, we performed a head-to-head comparison of high-dose intravenous (i.v.) and intratracheal (i.t.) vaccination in *Rag2^-/-^Il2rg^-/-^* double knockout mice, a severely immunodeficient model lacking T, B, and NK cells (23). To determine if pulmonary delivery improves survival, and limits systemic bacterial dissemination, mice received 10⁷ CFU of BCG, PE25SS, or BCG::RD1 either i.v. or i.t., a 10-20× higher dose than the standard vaccination dose of 0.5-1×10^6^ CFUs **(Fig. 1a, b)**.

**Figure 1:**
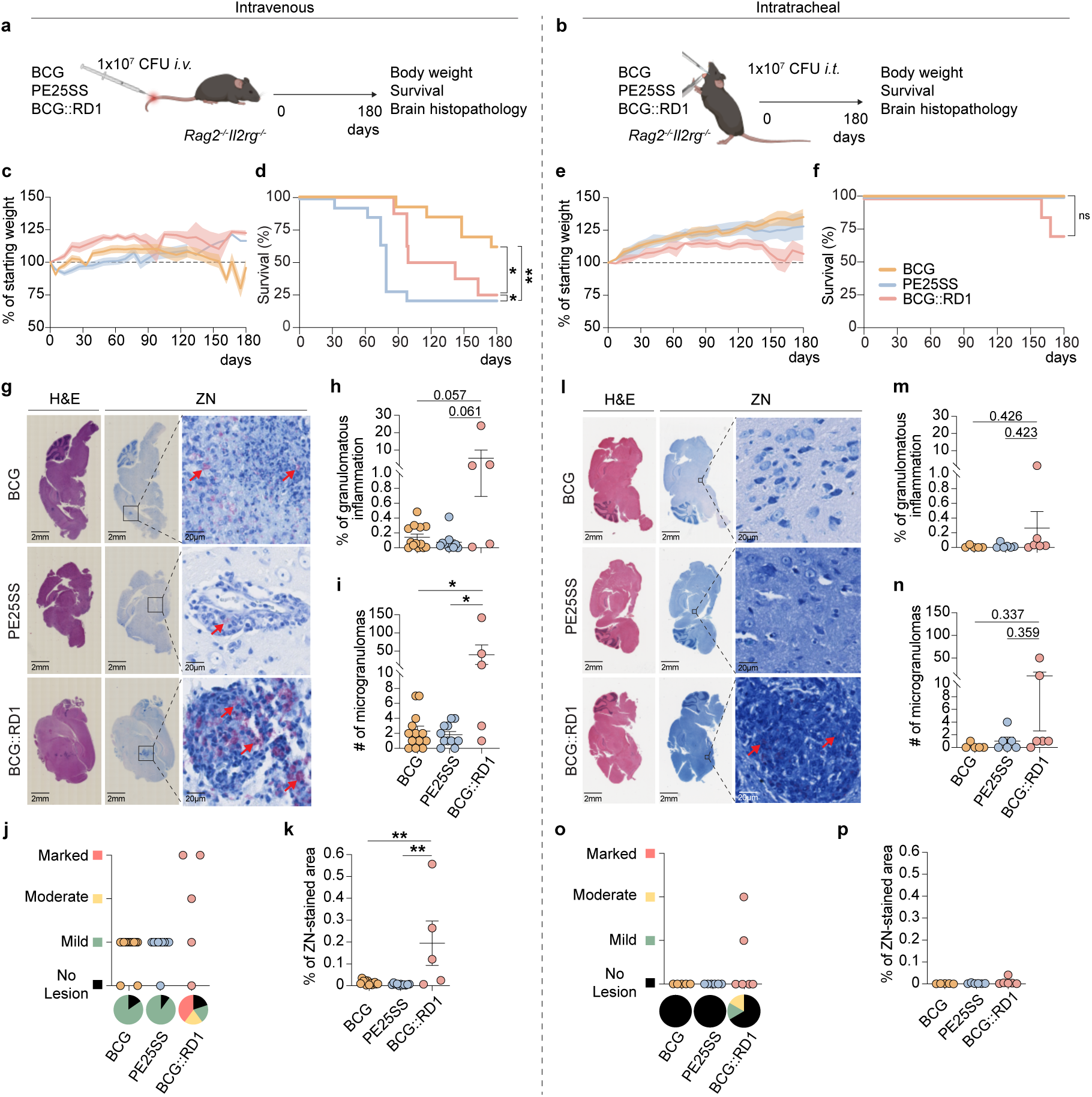
Pulmonary delivery overcomes the historical safety liabilities of ESAT-6-expressing rBCG in intravenous safety models. **(a)** *Rag2^-/-^Il2rg^-/-^* mice were intravenously and (**b)** intratracheally vaccinated with a high dose (10⁷ CFU) of BCG, PE25SS, or BCG::RD1 and monitored for 180 days. **(c, e)** Changes in body weight and **(d, f)** survival were monitored. **(g, l)** Representative images of H&E and ZN-stained brain sections. **(h, m)** The quantification of the percentage of granulomatous inflammation and **(i, n)** the total number of microgranulomas present in the brain. **(j, o)** Severity of pathological brain lesions was assessed and **(k, p)** the presence of mycobacteria was evaluated using ZN staining. Statistical significance was calculated using **(d, f)** Log-rank (Mantel-Cox) test or **(h, i, k, m, n, p)** One-way ANOVA with Tukey posthoc test (\**p* < 0.05, \*\**p* < 0.01, \*\*\**p* < 0.001).

High-dose i.v. delivery induced a transient loss of body weight in *Rag2^⁻/⁻^Il2rg^⁻/⁻^* mice, with all vaccine groups showing partial recovery approximately one-month later **(Fig. 1c)**. However, mice receiving parental BCG began to progressively lose weight after ∼2 months and ultimately exhibited the lowest average body weight at the 180-day endpoint. Among the rBCG strains, mice that received PE25SS succumbed earliest, with mortality beginning around 30 days, and displayed the lowest overall survival, closely followed by the BCG::RD1 group **(Fig. 1d)**. In contrast, i.v. parental BCG mice showed significantly greater long-term survival compared with both ESAT-6-expressing rBCG strains **(Fig. 1d)**. These findings are consistent with previous observations that high-dose systemic administration of antigen-enhanced rBCG strains can become detrimental in immunocompromised hosts. In contrast, all mice tolerated i.t. delivery remarkably well over the 180-day observation period **(Fig. 1e, f)**. All groups continued to gain weight until approximately day 120, after which only BCG::RD1-vaccinated mice exhibited a gradual decline in body weight, whereas BCG- and PE25SS-vaccinated mice continued to gain weight (**Fig. 1e**). Strikingly, both the parental BCG and PE25SS groups showed 100% survival at day 180, while only 2 of 7 BCG::RD1-vaccinated mice succumbed late in the study (days 160 and 168), resulting in a non-significant reduction in overall survival relative to the other groups (**Fig. 1f**).

In clinical settings, standard BCG vaccination in HIV-positive infants has been associated with disseminated BCGiosis (24, 25). Likewise, tuberculous meningoencephalitis (TBM) represents a severe manifestation of disseminated TB in immunocompromised individuals, particularly in people living with HIV and those with inherited immune defects (26), and remains associated with poor outcomes despite antibiotic treatment (27, 28). To assess mycobacterial dissemination and brain pathology, we quantified the extent of granulomatous inflammation in the H&E-stained brain **(Fig. 1g)**. Both i.v. BCG and PE25SS exhibited low levels of granulomatous inflammation, whereas i.v. BCG::RD1 induced substantially higher inflammation levels **(Fig. 1h)**. Similarly, i.v. BCG::RD1-vaccinated mice had a significantly greater number of microgranulomas than BCG and PE25SS, suggesting that RD1 expression is associated with increased brain pathology **(Fig. 1i)**. To further characterise brain involvement, brain lesions were categorised into four severity groups by a veterinary pathologist: no lesion, mild lesion, moderate lesion, and marked lesion **(Fig. 1j)**. Most i.v. BCG and PE25SS-vaccinated mice exhibited only mild lesions, whereas BCG::RD1-vaccinated mice developed both moderate and marked lesions. Additionally, ZN staining indicated significantly higher bacterial presence in i.v. BCG::RD1-infected brain sections, whereas BCG and PE25SS groups had lower bacterial presence (**Fig. 1g, k)**. As seen in the survival data, histopathological examination of the brain further highlighted the marked impact of administration route on safety outcomes **(Fig. 1g, l)**. All three i.t. groups exhibited either no or minimal granulomatous brain inflammation, with the exception of one i.t. RD1 mouse; however, this did not reach statistical significance **(Fig. 1m)**. Similarly, the number of microgranulomas was statistically comparable across groups, despite two i.t. RD1 mice displaying higher granuloma counts, one of which was euthanized prior to the endpoint after becoming moribund (day 160) **(Fig. 1n)**. No inflammatory lesions were detected in the brains of i.t. BCG- or PE25SS-vaccinated mice, and the majority of BCG::RD1-vaccinated mice were similarly unaffected, with only two mice displaying mild-to-moderate lesions (**Fig. 1o**). Notably, no severe pathology was observed, in contrast to pronounced lesions seen following high-dose i.v. BCG::RD1 administration **(Fig. 1j)**. Similarly, lower bacterial burden was observed in the brains of mucosally vaccinated animals (i.e., in BCG::RD1 i.t. mean % of ZN-area 0.0095% vs i.v. mean % of ZN-area 0.1947%) **(Fig. 1p)**.

Collectively, these data demonstrate that even under conditions of extreme immunodeficiency, high-dose mucosal delivery substantially mitigates the systemic dissemination and neuropathology historically associated with ESAT-6-expressing rBCG strains, underscoring the critical influence of delivery route on the safety profile of live attenuated TB vaccine candidates.

### Pulmonary rBCG vaccination enhances bone marrow myelopoiesis and augments macrophage bactericidal capacity in the absence of adaptive immunity

Next, we sought to elucidate the underlying immune mechanisms driving this striking observation. Given that more severe forms of immunodeficiencies such as HIV or primary immunodeficiency syndromes primarily impair adaptive immunity, particularly T-cell responses (29, 30), we reasoned that the improved survival observed following mucosal rBCG in *Rag2^-/-^Il2rg^-/-^*mice likely reflects compensatory innate immune activation capable of substituting for absent adaptive immune control. To this end, we employed *Rag1^-/-^* and *Rag2^-/-^Ilrg^-/-^* mice to assess the innate immune responses elicited by mucosal vaccination in the absence of adaptive immunity **(Fig. 2a, i)**. Using a flow cytometry-based cellular phenotyping approach, we tested whether rBCGs can quantitatively enhance innate cell populations in the respiratory tract and influence the bone marrow. Thirty days following pulmonary delivery, when previous experiments demonstrated pronounced cellular augmentation (31), blood, spleen lung tissue, and bronchoalveolar lavage fluid (BALF) were collected for immunophenotyping of innate immune cell subsets **(Fig. 2a, i, Fig. S1a-c)**. Additionally, lung sections were histologically examined to evaluate the degree of cellular infiltration following mucosal vaccination.

**Figure 2:**
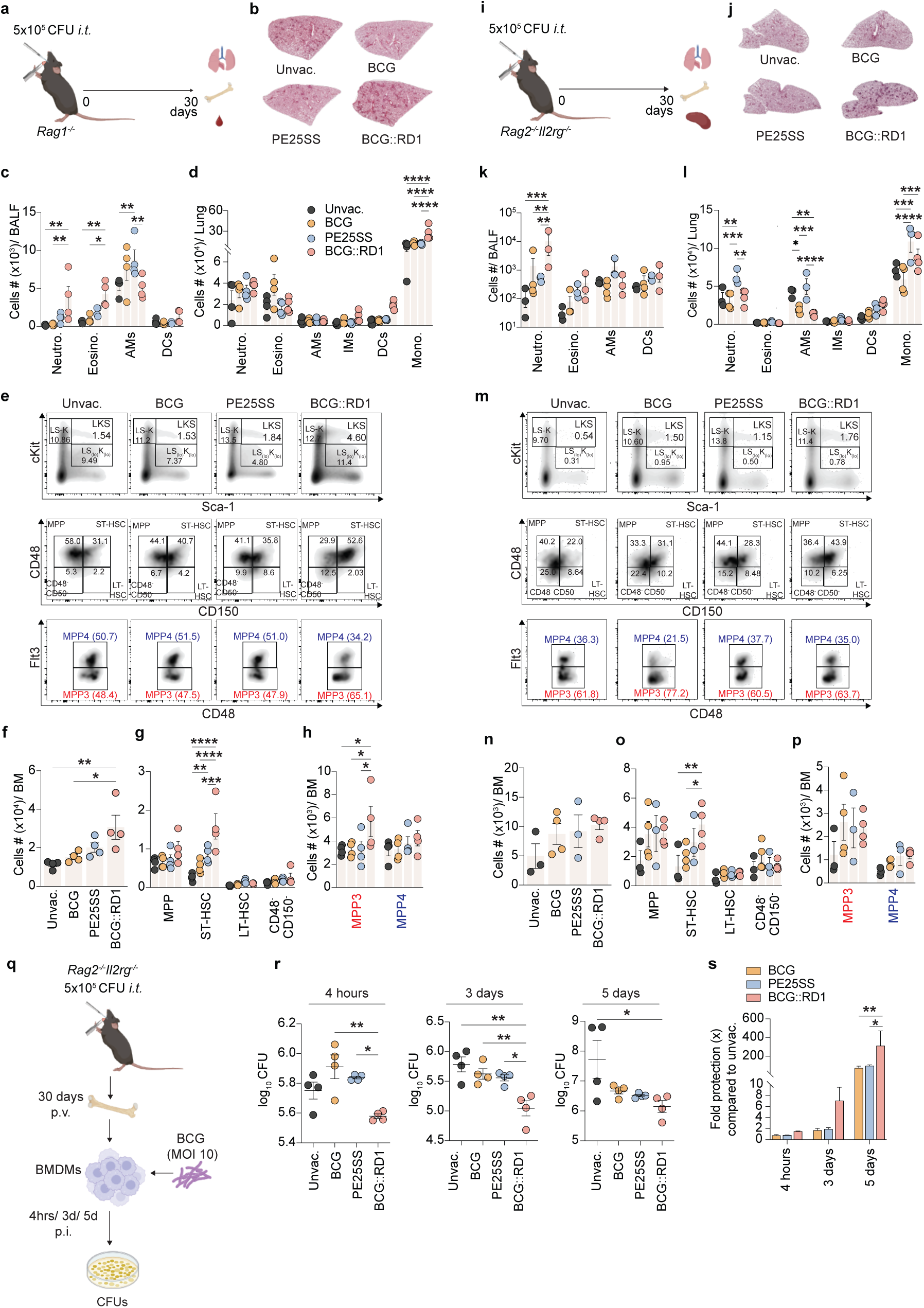
Pulmonary rBCG vaccination enhances bone marrow myelopoiesis and augment macrophage bactericidal capacity in the absence of adaptive immunity. FACS-based immune phenotyping was performed 30 days p.v. on cells isolated from BALF, blood, lung tissue and bone marrow of (a) *Rag1^-/-^* and (b) *Rag2^-/-^Il2rg^-/-^*mice. Representative images of H&E-stained lung sections from vaccinated (b) *Rag1^-/-^* and (j) *Rag2^-/-^Il2rg^-/-^* mice. **(c)** BALF and **(d)** lung-derived innate immune cells from *Rag1^-/-^* mice. **(k)** BALF and **(l)** lung-derived innate immune cells from *Rag2^-/-^Il2rg^-/-^* mice. Representative FACS plots of bone marrow Lin-cells from (e) *Rag1^-/-^* and (m) *Rag2^-/-^Il2rg^-/-^*mice. **(f)** Total and **(g)** subpopulations of LKS+ cells and **(h)** MPP3 vs MPP4 cells from *Rag1^-/-^*mice. **(n)** Total and **(o)** subpopulations of LKS+ cells and **(p)** MPP3 vs MPP4 cells from *Rag2^-/-^Il2rg^-/-^* mice. **(q)** BMDMs derived from *Rag2^-/-^Il2rg^-/-^* mice 30 days p.v. were assessed for bactericidal capacity using BCG as a surrogate mycobacterial challenge. **(r)** Viable bacteria count at 4 hrs, 3 days and 5 days. **(s)** Fold protection compared to unvaccinated group at 4 hrs, 3 days and 5 days. Statistical significance was calculated using **(f-h, n-p, r, s)** One-way ANOVA with Tukey posthoc or **(c, d, k, l)** Two-way ANOVA with Tukey posthoc test (\**p* < 0.05, \*\**p* < 0.01, \*\*\**p* < 0.001).

Overall cellular infiltration in these adaptive-immunity deficient lungs appeared largely comparable to that of unvaccinated lungs with slightly higher infiltration observed in the BCG::RD1 group **(Fig. 2b, j** and **Fig. 2c)**. Notably, in *Rag1^-/-^* mice, only the BCG::RD1 group showed a statistically significant increase in number of innate immune cells populations across BALF, lung and blood compared to the other vaccine groups **(Fig. 2c, d**, **Fig S2a, b)**. This includes marked expansion of BALF neutrophils, total lung monocytes and CD11b⁺ dendritic cells (DCs) **(Fig. 2c, d**, **Fig. S2b),** indicating that mucosal administration augments the local immune milieu at the primary site of vaccine delivery.

Prior studies have demonstrated that i.v. BCG can educate bone marrow stem cells, leading to enhanced myelopoiesis and the establishment of trained immunity (31). To determine whether mucosal vaccination, as shown in non-human primate models (32), induces trained immunity at the level of bone marrow haematopoiesis, we immunophenotyped lineage-negative (Lin⁻) enriched bone marrow cells from the vaccinated and unvaccinated mice **(Fig. S1d)**. Mucosal BCG::RD1 led to a significant increase in bone marrow progenitor populations in *Rag1^-/-^* mice **(Fig. 2e-h)**. Specifically, there was a marked expansion of the Sca-1⁺ c-Kit⁺ (LKS⁺) population, representing hematopoietic stem and progenitor cells (HSPCs) **(Fig. 2f)**. To further delineate these changes, we characterized subpopulations within the LKS⁺ compartment by measuring the proportions of long-term hematopoietic stem cells (LT-HSCs; LKS⁺CD150⁺CD48⁻), short-term HSCs (ST-HSCs; LKS⁺CD150⁺CD48⁺), and multipotent progenitors (MPPs; LKS⁺CD150⁻CD48⁺) **(Fig. 2e, g)**. A significant expansion of ST-HSCs was observed in the BCG::RD1 group compared to both standard BCG and PE25SS **(Fig. 2g)**. Further analysis focused on the polarization of MPP into myeloid versus lymphoid lineages. We assessed the balance between MPP3 (myeloid-biased) and MPP4 (lymphoid-biased) subsets. Although the relative proportions of myeloid-biased (MPP3) versus lymphoid-biased (MPP4) multipotent progenitors were comparable across all groups **(Fig. S2c)**, the total number of MPP3 cells was significantly higher in the BCG::RD1 group **(Fig. 2h)**, indicating a preferential expansion of bone marrow progenitor cells that are predisposed to differentiate into myeloid cells (33). This bias toward myelopoiesis is consistent with the potential induction of trained immunity, where enhanced production of myeloid cells (such as monocytes and neutrophils) plays a crucial role in improving anti-mycobacterial responses (31).

In *Rag2^-/-^Il2rg^-/^* mice, BCG::RD1 significantly increased the number of neutrophils in both the BALF and spleen, as well as total splenic monocytes, including classical monocytes **(Fig. 2k, Fig. S2d)**. Interestingly, while PE25SS led to a broader expansion of most lung innate immune populations in these mice **(Fig. 2l)**, BCG::RD1 still induced a pronounced increase in total and CD11b⁺ dendritic cells, and monocytes, including classical monocytes **(Fig. 2l, Fig. S2e)**. Although not reaching statistical significance, mucosal BCG::RD1 also increased LKS^+^ cells, particularly ST-HSCs and MPP3 in the bone marrow of *Rag2^-/-^Il2rg^-/-^* mice **(Fig. 2m-p, Fig. S2f)**, mirroring the pattern observed in *Rag1^-/-^* mice **(Fig. 2e-h)**.

Given the critical role of bone marrow-derived macrophages (BMDMs) in TB immunity (34), we next examined whether mucosal vaccination-induced alterations in HSCs and MPPs could influence the antimicrobial capacity of macrophages in the absence of adaptive immunity. To this end, BMDMs were generated from *Rag2^-/-^Il2rg^-/-^* mice 30 days after mucosal vaccination with BCG, PE25SS, or BCG::RD1, and their protective capacity was evaluated at 4 hours, 3 days, and 5 days post-infection (p.i.) using BCG as a surrogate mycobacterial challenge **(Fig. 2q)**. BMDMs from vaccinated *Rag2^-/-^Il2rg^-/-^* mice demonstrated better protection across all time points assessed **(Fig. 2r, s),** with the most pronounced bacterial reduction observed at 3 and 5 days p.i. suggesting that mucosal BCG::RD1 induces superior innate training compared to standard BCG.

Collectively, these findings indicate that ESAT-6-secreting BCG strains, particularly BCG::RD1, strongly activate local innate immunity at the respiratory interface of adaptive-deficient mice, marked by increased monocyte recruitment. The concurrent expansion of bone marrow progenitors suggests that BCG::RD1 retains a conserved capacity to promote myelopoiesis and innate inflammation even under severe immunodeficiency. Furthermore, the enhanced bactericidal activity of BMDMs from BCG::RD1-vaccinated *Rag2^-/-^Il2rg^-/-^* mice supports the notion that innate training contributes to the observed protection, enabling macrophages to mount a more effective antimicrobial response despite the absence of adaptive immunity. Notably, while i.v. administration of full-length RD1-expressing *Mtb* fail to induce bone marrow myelopoiesis in immunocompetent hosts with BCG::RD1 potentially exhibiting similar effects to *Mtb* (35), they appear to overcome this limitation in the absence of adaptive immunity, suggestive of two distinct mechanisms by which these strains influence the bone marrow, at least when administered mucosally.

### Pulmonary rBCG induces distinct transcriptional changes, but RD1-driven bone marrow myelopoiesis is weaker in immunocompetent mice

To further delineate the impact of distinct ESAT-6 secretion mechanisms (ESX-1 vs. ESX-5) and pulmonary delivery on lung and bone marrow, we extended our investigations into immunocompetent hosts. To this end, 30 days after i.t. delivery of vaccines into C57BL/6 mice, organs were collected for immunophenotyping of innate immune cell subsets **(Fig. 3a)**. Compared to adaptive-deficient *Rag1^-/-^* and *Rag2^-/-^Ilrg^-/-^*mice **(Fig. 2b, j)**, all vaccinated C57BL/6 mice showed notably increased cellular infiltration in lungs, indicating the presence of adaptive immune cells, with the highest infiltration observed in the rBCG groups **(Fig. 3b)**. All vaccines led to a marked expansion of one or more innate cell populations, including neutrophils, monocytes and DCs across the organs tested **(Fig. 3c, Fig. S3a, b)**. Mice that received mucosal PE25SS showed a statistically significant increase in neutrophils and DCs in the BALF compared to other vaccine candidates **(Fig. 3c)**.

**Figure 3:**
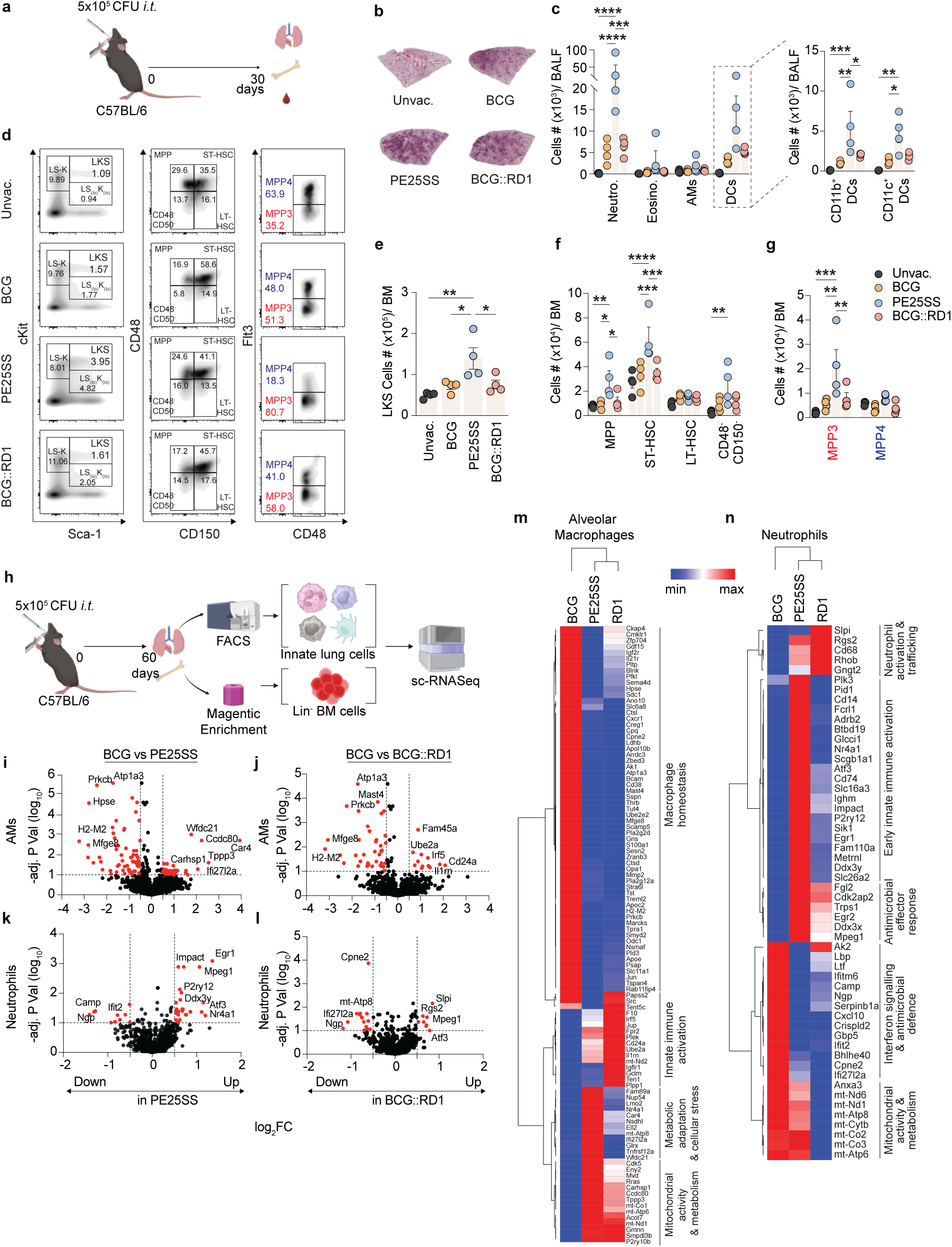
Mucosal rBCG induces distinct transcriptional changes but RD1-driven bone marrow myelopoiesis is weaker in immunocompetent mice. **(a)** Flow cytometry-based immune phenotyping was performed 30 days p.v. on cells isolated from BALF, blood, lung and bone marrow of C57BL/6 mice. **(b)** Representative images of H&E-stained lung sections from vaccinated animals. **(c)** BALF-derived innate immune cells. **(d)** Representative FACS plots of bone marrow Lin^-^ cells. **(e)** Total and **(f)** subpopulations of LKS^+^ cells and **(g)** MPP3 and MPP4 cells. **(h)** sc-RNASeq of FACS-sorted lung innate cells and magnetically enriched BM Lin^-^cells. Differentially expressed genes (DEGs; *false discovery rate [FDR] < 0.1, log_2_ fold change [Fc]>0.5*) of **(i, j)** AMs and **(k, l)** neutrophils compared to BCG group. Heatmaps showing normalised expression levels of DEGs in **(m)** AMs and **(n)** lung neutrophils. Statistical significance was calculated using **(e)** One-way ANOVA with Tukey posthoc test or **(b, f, g)** Two-way ANOVA with Tukey posthoc test (\**p* < 0.05, \*\**p* < 0.01, \*\*\**p* < 0.001).

Flow cytometry analysis revealed a notable expansion of the bone marrow LKS⁺ population, with the PE25SS group showing a statistically significant increase **(Fig. 3d, e)**. Further phenotyping revealed that although the overall expansion of LT-HSC, ST-HSCs and MPP populations was less profound than what has been observed following i.v. BCG administration in mice (31, 35), the rBCG groups, particularly the PE25SS group exhibited statistically significant increases in ST-HSCs and MPPs **(Fig. 3f)**. Remarkably, all vaccine groups, although most profound in the PE25SS group, showed a significant increase in the myeloid-biased MPP3 population relative to unvaccinated mice **(Fig. 3g, Fig. S3c)**. These phenotyping data suggest that pulmonary vaccination, particularly with PE25SS, enhances both local and systemic innate cell populations and influences the bone marrow, collectively contributing to an overall heightened innate immune state in immunocompetent mice. Interestingly, the lack of significant impact on the bone marrow by BCG::RD1 in C57BL/6 mice aligns with previous studies suggesting that *Mtb*, in fact, suppresses myelopoiesis in an RD1-dependent manner (35).

To further understand the impact of rBCG vaccination on lung cells, we performed single-cell RNA sequencing (scRNA-seq) to generate high-resolution transcriptional profiles of four key innate immune cell populations: alveolar macrophages (AMs; MerTK⁺ CD11c⁺ SiglecF⁺), interstitial macrophages (IMs; MerTK⁺ SiglecF⁻), DCs (CD11c⁺ MHCII⁺), and neutrophils (CD11b⁺ Ly6G⁺) isolated from the lungs, along with magnetically enriched Lin⁻ cells from the bone marrow of mice following 60 days of immune priming after mucosal delivery **(Fig. 3h, Fig. S1e, f** & **Fig. S3d)**. Among the cell types analysed, significant differential gene expression was observed only in AMs and neutrophils from rBCG-vaccinated lungs compared to standard BCG (**Fig. 3i-l, S3e, f**). In contrast, no notable transcriptional differences were detected in bone marrow HSPCs (**Fig. S3g**), possibly due to their unstimulated state prior to sequencing. AMs from rBCG vaccinated mice displayed distinct yet partially overlapping transcriptional profiles compared to standard BCG **(Fig. 3i, j, m)**. Higher expression of genes such as *Irf5* in BCG::RD1 lungs point to a more inflammatory phenotype (36), while PE25SS upregulated mitochondrial and metabolic genes (*mt-Atp6*, *mt-Co1*, *Nsdhl*, *Cdk5*), indicating a shift towards metabolic reprogramming **(Fig. 3m)**. PE25SS neutrophils exhibited a broader transcriptional shift, with upregulation of genes involved in metabolic adaptation (*Slc16a3*), innate immune activation (*Egr1*, *Cd14*), and immune regulation (*Nr4a1*) (37, 38), indicative of a transcriptionally activated state compared to both BCG and BCG::RD1 **(Fig. 3k, n).** These transcriptional changes in PE25SS are consistent with the significantly increased neutrophil expansion observed following PE25SS exposure **(Fig. 3c, n)**. Overall, rBCG, particularly PE25SS induced enhanced transcriptional reprogramming compared to parental BCG likely reflecting enhanced activation of innate myeloid cells at the lung of immunocompetent mice **(Fig. 3m, n**, **Fig. S3h, i)**. However, the reduced transcriptional response in lung neutrophils compared to PE25SS and absence of bone marrow myelopoiesis, support previous studies showing that RD1-containing strains (*Mtb* and BCG::RD1) regardless of delivery route fail to induce myelopoiesis in immunocompetent mice (35).

### Mucosally-induced innate priming confers protection against *Mtb* challenge across a spectrum of immunocompromisation

We next determined whether the enhanced *in vitro* killing capacity of BMDMs from immunocompromised mice also translates into protection against *Mtb* challenge across progressively immunocompromised hosts, using T2D, *Rag1^-/-^* and *Rag2^-/-^Il2rg^-/-^*mice. To address this, we first vaccinated diet-induced T2D (39) and age-matched non-diabetic control mice **(Fig. S4a-c)** i.t. with 5×10⁵ CFU of parental BCG, PE25SS or BCG::RD1, with a saline group serving as unvaccinated controls **(Fig. 4a)**. Sixty days post-vaccination (p.v.), all mice were challenged via aerosol with a low dose of *Mtb* (10-50 CFU), and 45 days later, lungs and spleen along with blood were collected to assess bacterial burden, histopathology, and systemic immune responses.

**Figure 4:**
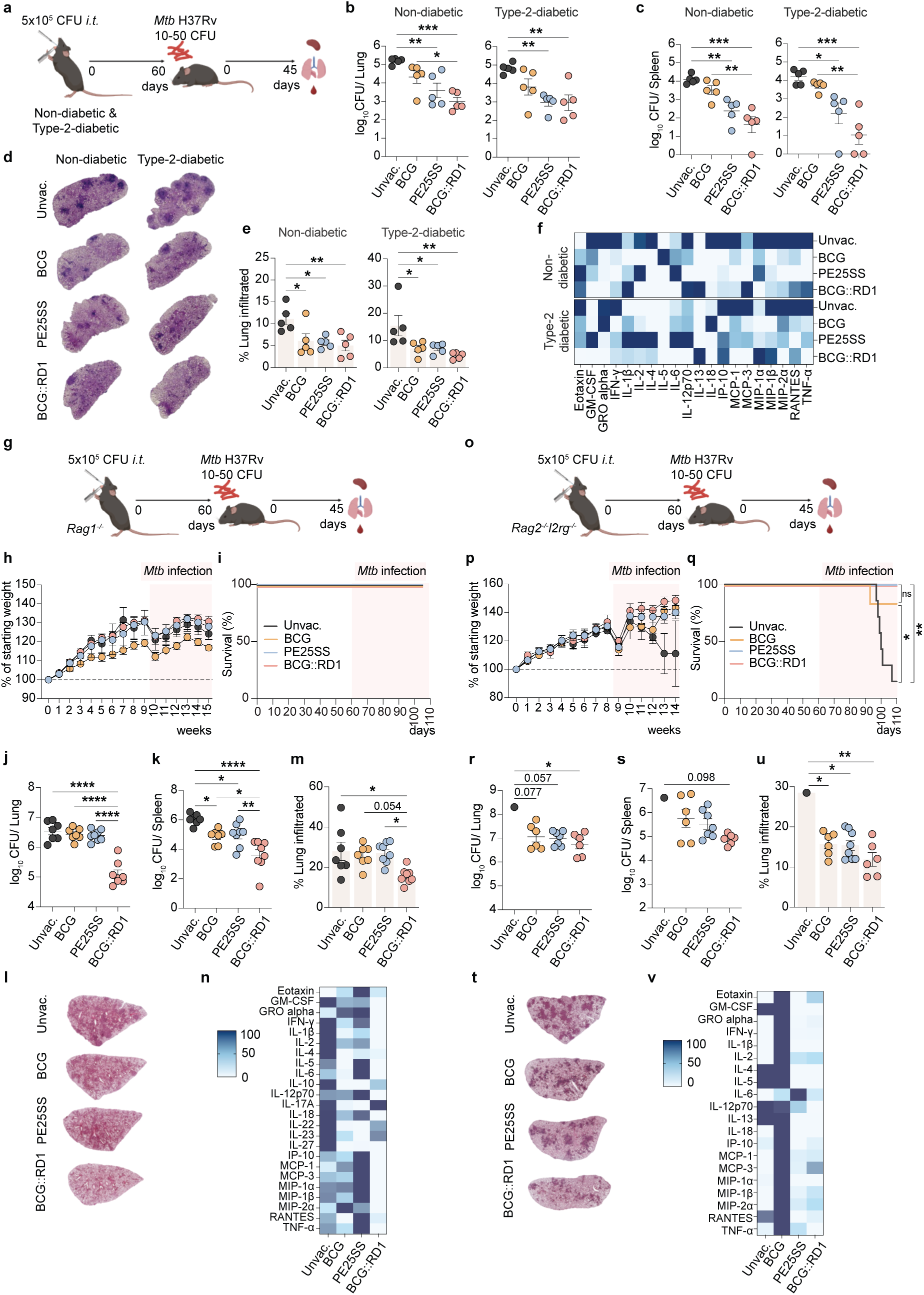
Pulmonary rBCG induced innate priming confers protection against *Mtb* across immunocompromised hosts. Schematic representation of vaccination and challenge experiment in **(a)** diabetic and non-diabetic mice, **(g)** *Rag1^-/-^*mice, and **(o)** *Rag2^-/-^Il2rg^-/-^*mice. **(b)** Lung and **(c)** spleen organ bacterial loads of diabetic and non-diabetic mice. **(d)** Representative images of H&E-stained lung sections and **(e)** quantitative assessment of lung tissue inflammation in diabetic and non-diabetic mice. **(f)** Heatmap showing normalized systemic cytokine and chemokine concentrations across experimental groups in diabetic and non-diabetic mice. **(h)** Body weight and **(i)** survival of *Rag1^-/-^* mice, and **(p)** body weight and **(q)** survival of *Rag2^-/-^Il2rg^-/-^* mice over the experimental duration. **(j)** Lung and **(k)** spleen organ bacterial loads of *Rag1^-/-^*, and **(r)** lung and **(s)** spleen organ bacterial loads of *Rag2^-/-^Il2rg^-/-^*mice, 45 days p.i.. Representative images of H&E-stained lung sections from **(l)** *Rag1^-/-^* and **(t)** *Rag2^-/-^Il2rg^-/-^* mice. Quantitative assessment of lung tissue inflammation in **(m)** *Rag1^-/-^* and **(u)** *Rag2^-/-^Il2rg^-/-^* mice following aerosol *Mtb* infection. Heatmap showing normalized z-score values of systemic cytokine and chemokine concentrations across experimental groups in **(n)** *Rag1^-/-^* and **(v)** *Rag2^-/-^Il2rg^-/-^* mice. Statistical significance was calculated using **(b, c, e, j-m, r-u)** One-way ANOVA with Tukey posthoc test or **(i, q)** Log-rank (Mantel-Cox) test (\**p* < 0.05, \*\**p* < 0.01, \*\*\**p* < 0.001).

Compared to unvaccinated controls, all vaccinated animals showed significantly lower *Mtb* burdens in both the lungs and spleens **(Fig. 4b, c)**. Notably, both BCG::RD1 and PE25SS groups harboured lower bacterial burdens than parental BCG, with similar patterns observed in the diabetic and non-diabetic cohorts. BCG::RD1 exhibited the lowest bacterial counts overall **(Fig. 4b, c)**. Lung histopathology further supported these findings **(Fig. 4d)**. Both non-diabetic and diabetic animals exhibited similar patterns of tissue inflammation; however, as expected, unvaccinated mice showed the most severe lung tissue damage **(Fig. 4d, e)**. Unvaccinated diabetic mice had greater inflammation than their non-diabetic counterparts, suggesting a more severe disease progression in diabetic animals (mean±SEM;11.08±1.319% vs. 15.45±3.754%) **(Fig. 4e)**. In contrast, all three vaccine groups significantly reduced inflammatory pathology **(Fig. 4e)**. The BCG::RD1 group showed the lowest levels of inflammatory cell infiltration, which is consistent with its ability to more effectively limit bacterial dissemination and tissue injury. Consistently, unvaccinated mice, both diabetic and non-diabetic, exhibited the highest overall average levels of pro-inflammatory serum cytokines and chemokines, suggestive of an unresolved infection **(Fig. 4f, Fig S4c-h)**. This confirms our previous findings that mucosal rBCGs, particularly BCG::RD1, confer superior protection against *Mtb* compared with parental BCG, including in T2D comorbid immunodeficient hosts.

To determine whether these rBCG strains could also protect more severely immunocompromised hosts in a manner completely independent of adaptive immunity, we first challenged i.t. rBCG-vaccinated *Rag1^-/-^* mice with a low dose of *Mtb* **(Fig. 4g)**. Throughout the study period, all *Rag1^-/-^* mice gained weight, except for a reduction during the first week following *Mtb* infection; however, the animals recovered and reached a peak body weight at 13 weeks p.v., followed by a gradual decline until the end of the experiment **(Fig. 4h)**. Notably, the overall body weight patterns were similar across all groups, and 100% survival was observed following both vaccination and infection, indicating that pulmonary administration of all vaccines was well tolerated even in the absence of adaptive immunity **(Fig. 4i)**. When organ bacterial loads were determined at 45 days p.i., unvaccinated *Rag1^-/-^*mice harboured approximately ∼6.5 and ∼6 log_10_ *Mtb* CFUs in their lungs and spleens, respectively, underscoring their extreme susceptibility to *Mtb* due to the absence of lymphocytes **(Fig. 4j, k)**. In contrast, all vaccinated groups exhibited reduced *Mtb* CFUs, with the BCG::RD1 group showing a significantly lower bacterial burden of nearly 2 log_10_ in both the lungs and spleen compared to unvaccinated controls as well as to the standard BCG and PE25SS groups **(Fig. 4j, k)**. In the spleen, both standard BCG and PE25SS also significantly reduced CFUs compared to the unvaccinated group **(Fig. 4k)**. This reduction in bacterial load was mirrored by lung histopathology, where lung sections from BCG::RD1-vaccinated mice displayed the least inflammatory cell infiltration and tissue damage **(Fig. 4l, m)**. Systemic cytokine and chemokine analyses revealed that unvaccinated *Rag1^-/-^*mice had the highest levels of pro-inflammatory mediators such as IFN-γ, TNF-α, IL-12 and IL-18, indicative of unresolved infection, whereas mice vaccinated with BCG::RD1 exhibited the lowest cytokine levels **(Fig. 4n, Fig. S5a-f)**, correlating with their reduced bacterial burden and improved lung pathology **(Fig. S5g)**. This was further supported by a PCA analysis, which integrated serum cytokine profiles, lung pathology and organ bacterial burdens, where BCG::RD1-vaccinated mice clustered distinctly from other groups, reflecting their unique inflammatory and protective profile **(Fig. S5h)**. IL-12p70 and IL-6 levels were increased in some of the PE25SS mice **(Fig S5d, f)**.

We next employed *Rag2^-/-^γc^-/-^* mice to further evaluate the efficacy and safety of these vaccines in hosts entirely lacking adaptive immunity, including NK cells (**Fig. 4o)**. Surprisingly, despite their severe immunodeficiency, all mice gained weight (**Fig. 4p)** and survived the 60-day mucosal vaccination phase (**Fig. 4q)**. While this was in stark contrast to classical i.v. infection in *Rag2^⁻/⁻^Il2rg^⁻/⁻^* mice **(Fig. 1d)** (40), it was consistent with our earlier observation that *Rag2^⁻/⁻^Il2rg^⁻/⁻^* mice tolerated and survived high-dose mucosal administration of rBCGs **(Fig. 1f)**. Following aerosol *Mtb* challenge of i.t. vaccinated mice, a notable temporary drop in body weight was observed, from which all groups initially recovered within a week (**Fig. 4p)**. Subsequently, 6 out of 7 unvaccinated mice succumbed to the infection before the study endpoint (45 days p.i.), leaving only one surviving animal, which underscores the extreme susceptibility of this strain to *Mtb* in the absence of adaptive immunity **(Fig. 4q)**. In stark contrast, among the vaccinated groups, only one mouse in the parental BCG group reached ethical endpoints, while both groups receiving the ESAT-6-secreting vaccines (BCG::RD1 and PE25SS) exhibited 100% survival **(Fig. 4q)**. When organ bacterial loads were determined, the surviving unvaccinated mouse harboured approximately ∼8 log_10_ CFU in the lungs, although statistical comparisons to other groups were limited by the low sample size of one surviving mouse **(Fig. 4r)**. On the other hand, all vaccinated mice showed a notable reduction in lung CFU of more than 1 log, with the BCG::RD1 group achieving a statistically significant reduction of ∼1.5 log compared to the unvaccinated control **(Fig. 4r)**. In the spleen, while no statistically significant differences were observed among the groups due to low statistical power, the BCG::RD1 group again exhibited the lowest bacterial counts of about a 1.7 log reduction relative to unvaccinated mice, and a nearly 1 log reduction relative to BCG vaccinated mice **(Fig. 4s)**. Lung histopathology mirrored these findings: the unvaccinated mouse displayed the highest degree of inflammatory infiltration, whereas all vaccinated groups showed substantially reduced lung inflammation, with the BCG::RD1 group reaching the lowest inflammation levels **(Fig. 4t, u)**. When serum was assessed for cytokine and chemokine levels at the study endpoint, the BCG-vaccinated group exhibited the overall highest concentrations across most analytes compared to other vaccine groups **(Fig. 4v, Fig. S5i)**, with the caveat that serum for the unvaccinated group was only available from the one surviving mouse. Notably, levels of key pro-inflammatory cytokines, IFN-γ, TNF-α and IL-12p70 were significantly elevated in the BCG group relative to both BCG::RD1 and PE25SS groups **(Fig. S5j-l)**.

Overall, these data show that all vaccines were well tolerated across host backgrounds, from immunocompetent C57BL/6mice to severely adaptive immune-deficient *Rag1^-/^* and *Rag2^-/-^Il2rg^-/-^*mice. Notably and paradigm-changing, ESAT-6-secreting rBCG strains, particularly BCG::RD1, markedly reduced organ bacterial burden and lung inflammation, indicating that robust innate immune priming alone can confer significant protection against low-dose aerosolized *Mtb* infection, even in the absence of adaptive immunity. Collectively, these findings suggest that pulmonary delivery of rBCGs drives potent innate immune activation that may mitigate traditional safety concerns associated with *in vivo* ESAT-6 secretion virulence while preserving strong protective efficacy. It is particularly intriguing that the most virulent strain, BCG::RD1, conferred the greatest protection in the absence of adaptive immunity.

## Discussion

Secreted proteins of *Mtb* have long been known to be a rich source of immunogens (41). Experimental LAVs expressing immunodominant antigens, particularly from the *Mtb* RD1 region, have shown improved protection compared to BCG (14, 15). However, safety concerns linked to increased persistence and virulence have hindered their clinical translation (14, 15). As a result, more than 100 years on, BCG still remains the benchmark in protection against TB. In this study, we demonstrate that pulmonary delivery of BCG strains carrying the full-length RD1 region (BCG::RD1) or exporting ESAT-6 via the native ESX-5 secretion system (PE25SS) is not only (i) well tolerated but also (ii) confers superior protection compared to standard BCG. Remarkably, this enhanced safety and efficacy extend beyond immunocompetent hosts to preclinical models of more severe immunodeficiencies, highlighting the potential of these LAVs for broader and safer application in vulnerable populations.

The improved safety profiles of antigen-enhanced BCG strains observed in this study sharply contrast with outcomes from i.v. infections in SCID, *TNF-α^-/-^* and *Ifngr1*^-/-^ mice (12, 41), highlighting the critical importance of dose and delivery route in LAV safety assessments. Notably, even among licensed BCG strains, safety and survival outcomes can differ markedly in i.v. SCID infections (16), underscoring the complex, context-dependent nature of vaccine safety assessments. This raises a critical question; what is a physiologically relevant pre-clinical safety threshold for new live attenuated TB vaccine candidates? While commonly used i.v. models remain useful, they risk prematurely excluding promising candidates that may be both safe and effective when delivered via more clinically relevant routes such as pulmonary administration.

Mucosal delivery of LAVs, including standard BCG, has shown enhanced protective efficacy driven in part by the induction of T_RM_ in animal models, including in non-human primates (42, 43). Supporting this, a recent human aerosol BCG infection study demonstrated rapid recruitment of innate and BCG-specific adaptive immune cells to the lung mucosa (44). Mucosal vaccination can also promote local innate responses that can limit early bacterial replication and dissemination (45). In a previous study, we demonstrated that the superior protection conferred by mucosal delivery of BCG::RD1 in a TB/T2D comorbid model was associated with an enhanced innate immune response (21). scRNA-seq analysis in the current study demonstrated that mucosal vaccination with rBCGs reprogrammed the transcriptional landscape of lung tissue-resident macrophages and neutrophils compared to standard BCG, indicative of enhanced phagocytic function and antimicrobial activity. These transcriptional changes suggest that early innate control of *Mtb* is driven, at least in part, by the enhanced functional capacity of lung-resident macrophages and neutrophils primed by these ESAT-6-secreting LAVs.

In immunocompetent mice, i.v. *Mtb* failed to enhance myelopoiesis compared to the parental BCG (35), whereas in the current study, pulmonary PE25SS did promote increased myeloid differentiation at 30 days p.v., but not BCG::RD1. Given that both BCG::RD1 and PE25SS export ESAT-6, these findings suggest that other RD1-encoded proteins beyond ESAT-6 may play a detrimental role in the suppression of bone marrow myelopoiesis. It would therefore be particularly valuable to investigate the roles of RD1-encoded virulence factors such as Rv3865 and Rv3866 (46, 47), as well as ESAT-6 itself, in modulating bone marrow progenitor dynamics independently of ESAT-6 secretion. While i.v. *Mtb* fail to induce myelopoiesis due to suppression mediated through the type I interferon/iron (IFN-I/Fe) axis (35), mucosal delivery of ESAT-6 may instead prime local myeloid cells via an ESAT-6-CFP-10-dependent mechanism potentially involving phagosomal rupture and cytosolic access. This localized priming may have conferred enhanced protection against *Mtb* without relying heavily on bone marrow-driven myelopoiesis in the current context. In fact, this may be attributed to the synergistic interplay between the innate and adaptive immune systems in immunocompetent hosts, where adaptive responses may mask innate-only differences despite the enhanced protection.

In severely immunocompromised *Rag1^⁻/⁻^* mice, BCG::RD1 markedly expanded the bone marrow progenitor compartment and enhanced myelopoiesis compared to PE25SS, indicating that RD1-dependent suppression is alleviated in the absence of adaptive immunity. One potential explanation is that restoration of the RD1/ESX-1 secretion system in BCG::RD1 extends beyond ESAT-6 antigen delivery. ESX-1 has been shown to cooperate with PDIM lipids to disrupt the phagosomal membrane, allowing mycobacterial products to access the cytosol and activate innate immune sensing pathways (48, 49). Consistently, BMDMs from *Rag2^-/-^Il2rg^-/-^* mice vaccinated with ESAT-6-secreting BCG strains, particularly BCG::RD1, displayed superior *in vitro* bacterial killing, supporting the notion that these vaccines reprogram bone marrow HSPCs to generate functionally enhanced, trained myeloid progeny. In this context, the superior protection observed against aerosol *Mtb* infection is likely orchestrated by an enhanced innate response driven by increased recruitment and activation of monocytes and neutrophils that enables early bacterial control in the absence of adaptive immunity. These observations suggest that in the absence of adaptive immunity (as in *Rag1^⁻/⁻^* and *Rag2^-/-^Il2rg^-/-^*mice), the suppressive signals may be lost, likely due to altered cytokine milieu and disrupted feedback loops, permitting progenitor expansion and more robust innate cell replenishment. Furthermore, these findings collectively suggest the presence of two distinct mechanisms by which BCG::RD1 and PE25SS influence the bone marrow, particularly following mucosal administration, an observation that warrants further investigation to delineate the underlying pathways and their implications for vaccine-induced innate training.

Overall, our findings demonstrate that ESAT-6-expressing BCG strains are safe and provide superior protection compared to parental BCG when delivered directly into the murine lung. The inclusion of a broader antigenic repertoire, encompassing highly immunogenic *Mtb* antigens, enhances vaccine immunogenicity and reshapes the transcriptional landscape of myeloid cells, including macrophages, thereby contributing to improved protection against *Mtb* (50, 51). Moreover, in the absence of adaptive immunity, BCG strains carrying the full-length RD1 locus appear to promote the expansion of bone marrow progenitors, generating a functionally enhanced myeloid compartment capable of driving early control and clearance of *Mtb* infection. These findings highlight the potential of leveraging ESAT-6-based or RD1-containing BCG platforms, such as BCG::ESX-1^Mmar^, a rBCG strain which heterologously expresses the esx-1 region of *M. marinum* (52, 53), as a foundation for next-generation vaccines specifically tailored for immunocompromised hosts, in whom standard BCG may be either unsafe or insufficiently protective. In summary, we demonstrate vaccine delivery route as a critical determinant of safety and suggest that pulmonary administration may unlock the clinical potential of highly efficacious TB LAV candidates previously limited by safety concerns.

## Materials and Methods

### Animals

All animal experiments were approved by the Animal Ethics Committee (AEC) of James Cook University (A2400, A2794 and A2837). All mouse strains (C57BL/6, *Rag1^-/-^*, *Rag2^-/-^Il2rg^-/-^*) were bred and maintained in specific-pathogen free (SPF) animal facilities within the Australian Institute of Tropical Health and Medicine (AITHM), James Cook University, Australia. The murine T2D model employed in this study was developed and extensively characterized using male C57BL/6 mice (39). In brief, at 4-to 6-week-old male C57BL/6 mice were randomly assigned into one of two dietary groups for a 30-week intervention. One group had ad libitum access to an EDD (SF03-030, Specialty Feeds, Western Australia), a high glycaemic index, semi-pure rodent diet containing 23% fat, 19.4% protein, 50.5% dextrose, and 9.32% fibre, while the control group received an isometric quantity of SD (SF08-020, Standard AIN93M rodent diet) containing 4% fat, 13.8% protein, 64.8% carbohydrate, and 9.4% fibre). After 30 weeks, mice from both groups were evaluated for body weight gain, fasting blood glucose levels and glucose tolerance to determine the diabetic status (39). In other experiments, 6-to 8-week-old young adult C57BL/6, *Rag1^-/-^* and *Rag2^-/-^Il2rg^-/-^* mice were used.

### Bacterial strains

*Mycobacterium tuberculosis* strain H37Rv and all BCG strains (BCG Pasteur, BCG::RD1 and BCG::ESAT6-PE25SS) were cultured as previously described (54). Briefly, bacterial strains were expanded in 7H9 broth (BD Bioscience) supplemented with 0.2% w/v glycerol, 0.05% w/v, Tween 80, 10% v/v, Middlebrook albumin-dextrose-catalase (ADC) enrichment (BD Biosciences) and the appropriate antibiotics. Mid-logarithmic cultures were harvested, washed in sterile phosphate-buffered saline (PBS), and stored in 15% glycerol/PBS at −80 °C until required for vaccination or infection.

### Immunization and infection

Prior to vaccination and infection, frozen bacterial stocks were thawed, sonicated, centrifuged, and the resulting pellets were resuspended in sterile PBS to achieve the desired vaccination or infection dose. In all vaccine experiments, mice were immunized via intratracheal (i.t.) administration with 5×10^5^ CFUs in a total volume of 50 μl per mouse (21). For safety assessments, *Rag2^-/-^Il2rg^-/-^* mice were vaccinated i.t. or intravenously (i.v.) with 10^7^ CFUs in 100 μl per mouse. Sixty days p.v., mice were challenged with a low aerosol dose of *Mtb* H37Rv (10 to 50 CFUs) using a Glas-Col inhalation exposure system in a BSL3 (PC3) laboratory (55). The exact initial infectious dose was confirmed by plating homogenized lung tissue collected from mice 1-day p.i. onto Middlebrook 7H11 agar supplemented with 10% oleic acid-albumin-dextrose-catalase (OADC).

### Determination of organ bacterial loads

Lung and spleen tissues were homogenized in sterile sample bags or gentleMACS C tubes containing 1 mL of sterile PBS/0.05% Tween 80. Serial dilutions of the homogenates were plated 7H11 agar plates enriched with 10% OADC and supplemented with 10 µg/mL cycloheximide and appropriate antibiotics (50 µg/mL hygromycin and 20 µg/mL ampicillin for rBCG strains and *Mtb*, respectively). To restrict the growth of BCG strains during *Mtb* culture, Thiophene-2-carboxylic acid hydrazide (TCH, 2 µg/mL; Sigma) was added to 7H11 agar plates. The agar plates were then sealed and incubated aerobically at 37 °C for 4 to 5 weeks. Colonies were subsequently counted and the total CFUs per organ were calculated based on dilution factors and the organ size.

### Histology

Perfused or unperfused lung lobes and brains were fixed overnight in 4% paraformaldehyde and transferred into 70% ethanol the following day until further processing. The organs were then embedded in paraffin, sectioned at 4 µm, and stained with hematoxylin and eosin (H&E) and/or acid-fast Ziehl-Neelsen (ZN) stain. ImageJ and QuPath software were used to identify mycobacteria and quantify the percentage of tissue affected by dense cellular infiltration by measuring the total tissue area and the area of infiltrates, as described previously (21).

### Single cell suspension for flow cytometry and magnetic enrichment

Lungs were perfused with 20 mL of sterile PBS, excised, mechanically disrupted, and digested for 30 minutes at 37 °C in sterile RPMI 1640 medium supplemented with 10% heat-inactivated fetal bovine serum (FBS; Gibco), 100 U/mL penicillin, 100 µg/mL streptomycin (Gibco), 7.5 µg/mL Collagenase D (Sigma), 1.75 µg/mL Collagenase VIII (Sigma), and 200 µg/mL DNase I (Sigma). Single-cell suspensions were generated by passing the digested tissue through a 70-µm cell strainer, followed by erythrocyte lysis using ACK buffer (ref). Bone marrow cells were harvested from femurs and tibiae in cold, sterile RPMI. These cells were either directly subjected to antibody staining or, for isolation of untouched stem and progenitor cells, magnetically enriched using the EasySep Mouse Hematopoietic Progenitor Cell Isolation Kit (STEMCELL Technologies) **(Fig. S1f)**. Additionally, cell suspensions from bronchoalveolar lavage fluid (BALF) and 100 μL of whole blood were lysed with ACK buffer prior to staining.

### Flowcytometry

Immune cell phenotyping of blood, lung, BALF and bone marrow was done using the following antibodies. CD3e (500 A2), NKp46 (29A1.4), CD103 (M290), CD11c (HL3), CD11b (MI/70), F4/80 (BM8), I-A/I-E (M5/114.15.2), CD64a/b (X54-9/7.1), CD40 (3/23), CD80 (16-10A1), CD86 (GL-1), Ly6G (1A8), Siglec-F (E50-2440), Ly6C (AL-21) were obtained from BD Biosciences. CD103 (2E7), CD11c (N418) MerTK (DS5MMER), Ly-6A/E (Sca-1) (D7), CD117 (c-Kit), CD34 (RAM34), CD150 (mShad150), CD48 (HM48-1), CD135 (Flt3, A2F10), CD16/32 (93), CD127 (A7R34), Mouse Hematopoietic Lineage Antibody Cocktail (17A2, RA3-6B2, M1/70, TER-119, RB6-8C5) were purchased from eBioscience (ThermoFisher). Zombie UV Fixable viability kit (Biolegend) and Fixable viability stain 780 (BD Biosciences) were used to exclude dead cells. Cells were enumerated using CountBright Absolute Counting Beads (Invitrogen). Cell acquisition was performed on a BD Fortessa X20 using FACSDiva in FACS buffer (PBS with 5% FBS and 0.1% NaN₃), and data were analysed with FlowJo software version 10 (BD Biosciences). Where applicable, the same antibodies were used for FACS sorting of lung cells, which were sorted on a BD FACSAria III. Gating strategies for different immune cell populations are shown in **Fig. S1a-e**.

### Serum cytokine and chemokine analysis

To determine circulating cytokine and chemokine levels, blood was collected via cardiac puncture into Z-gel tubes (Sarstedt). Serum was separated and filtered through 0.2 µm SpinX columns (Sigma; for *Mtb*-infected animals) and stored at –20°C. Prior to analysis, frozen serum samples were thawed and prepared according to the mouse ProcartaPlex Th1/Th2/Chemokine 20- and 26-plex kit instructions (ThermoFisher). Samples were then analysed using a MagPix (Luminex) instrument along with the ProcartaPlex Analysis App (ThermoFisher).

### Single cell RNA sequencing and data analysis

Single-cell RNA-seq was performed using the 10x Genomics Chromium Next GEM Flex Gene Expression assay, following the manufacturer’s protocol (10x Genomics, CG000477, Rev D). Fixed, pooled, cell suspensions from FACS-sorted innate lung and magnetically enriched Lin-bone marrow were thawed, washed, and counted using AO-PI staining on a LUNA-FL Automated Cell Counter. Cells were then hybridised with mouse WTA probes for 23 hours at 42°C. Following hybridisation, cell concentrations were redetermined. GEM encapsulation was performed on the Chromium X instrument, targeting the maximum number of cells for capture per sample. GEMs were recovered and processed, and the resulting products were amplified and indexed to generate sequencing libraries. Library quality was assessed using the Agilent TapeStation D1000 platform and quantified by qPCR. Libraries were then pooled at equimolar concentrations and sequenced on the Illumina NovaSeq X Plus platform, following 10x Genomics recommended guidelines of at least 10,000 read pairs per cell. Each of the four samples were processed with cellRanger and imported as a singleCellObject into R for preprocessing, clustering and annotation following established pipelines (56–58). Doublets were called with SCDS and scDblFinder (59, 60), and damaged cells were identified based on the predicted mitochondrial content with those showing three standard deviations of mitochondrial content above the mean removed. Cells with likewise high ribosomal content were also removed. The singleCellObject was then converted to a Seurat object and cells lacking expression for more than 95% of all genes were also discarded (61). Five cell types were annotated from a total of 3,600 high quality single cells - alveolar macrophages (MerTK^+^ SiglecF^+^), non-alveolar macrophages (MerTK^+^ SiglecF^-^), dendritic cells (CD11c^+^ MHCII^+^), neutrophils (CD11b^+^ Ly6G^+^) and Kit^+^ Sca^+^ cells. Count normalisation was conducted with Sanity prior to differential expression analysis with Limma which followed the workflow described elsewhere (62–64). Unsupervised clustering and DEG (*false discovery rate [FDR] < 0.1, log_2_ fold change [Fc]>0.5*) expression data was visualised by Morpheus software (https://software.broadinstitute.org/morpheus).

### Generation of BMDMs

Bone marrow from both femurs and tibiae was harvested in cold RPMI 1640 supplemented with 10% heat-inactivated FBS (Gibco). Cells were subsequently seeded in DMEM supplemented with 10% FBS, 2mM L-glutamine, 1mM sodium pyruvate, 100U/ml penicillin, 100μg/ml streptomycin, and 50 ng/ml mouse M-CSF (Gibco) in 6-well plates. After 3 days of incubation at 37°C with 5% CO2, fresh medium containing M-CSF was added. Bone marrow cells were differentiated into macrophages over 6–7 days and examined microscopically to confirm adherence. Cells were subsequently harvested by removing the supernatant and incubating with 1 mL of pre-warmed TrypLE (Gibco) per well for 10 minutes at 37°C. Macrophage purity was assessed by flow cytometry based on surface expression of CD11b and F4/80 **(Fig. S1g)**.

### *In vitro* macrophage killing assay

*In vitro* macrophage killing assay was performed as previously described (31, 35). Briefly, BMDMs (∼3×10^5^ cells) were seeded in 12-well plates supplemented with DMEM without penicillin/streptomycin (500 μl), and incubated overnight at 37°C with 5% CO_2_. Next day, cells were infected with BCG (MOI 10), unless otherwise indicated. Cells were incubated for 4 hours at 37°C with 5% CO_2_. Subsequently, cells were washed 3x with sterile PBS, and were then incubated in supplemented DMEM without penicillin/streptomycin. CFUs were enumerated at 4 hours, day 3 and day 5 post-infection.

### Statistics

All statistical analysis was performed, and graphs were generated using Prism version 10 (GraphPad). For two-group comparisons, the Students’ t-test was used. For comparisons across multiple groups, one-way or two-way ANOVA with Tukey’s or Dunnett’s post-hoc test was applied, depending on data normality. Survival curves were analysed using Gehan-Wilcoxon test. Cytokine and chemokine data were z-score normalized for heatmap visualization. P < 0.05 was considered statistically significant.

## Data Availability

Requests for further information and resources should be directed to and will be fulfilled by the lead contact, Andreas Kupz. This study did not generate new, unique reagents. All data reported in this paper will be shared by the lead contact upon request.

## Code availability

This paper does not report original codes. Any additional information required to reanalyze the data reported in this paper is available in the main text and supplemental information or from the lead contact upon request.

## Acknowledgements

We thank Serrin Rowarth, Kylie Robertson, Sophie-Marie Gisder and Leanne Taylor for support in the animal facility and BSL3 laboratory.

## Author contributions

Conceptualisation: H.D.S, A.K. Investigation: H.D.S, G.Z, A.C, M.P, S.M.H, Y.W, J.S, A.K. Manuscript writing: H.D.S, A.K. Manuscript editing: H.D.S, G.Z, A.C, M.P, S.M.H, Y.W, J.S, M.A.F, A.K. Funding acquisition: A.K, M.A.F.

