## supplementary figure 1-5 for "Pulmonary delivery of antigen-enhanced BCG overcomes safety barriers in immunocompromised hosts and protects against TB in the absence of adaptive immunity"

**Supplemental Materials**

Harindra Darshana Sathkumara^1^, Guangzu Zhao^1^, Andrew Calcino^1, 2, 3^, Munish Puri^1^, Socorro Miranda-Hernandez^1^, Yide Wong^1^, Julia Seifert^1^, Matt A. Field^1, 2, 3^, Roland Brosch^4^, Andreas Kupz^1, *^

^1^Australian Institute of Tropical Health and Medicine, James Cook University, Cairns & Townsville, Queensland, Australia

^2^Centre for Tropical Bioinformatic and Molecular Biology, Australian Institute of Tropical Health and Medicine, James Cook University, Cairns, Queensland, Australia.

^3^College of Science and Engineering and Centre for Tropical Bioinformatics, James Cook University, Townsville, Queensland, Australia.

^4^Institut Pasteur, Université Paris Cité, Unit for Integrated Mycobacterial Pathogenomics, CNRS UMR 6047, Paris, France

Correspondence

*Prof Andreas Kupz

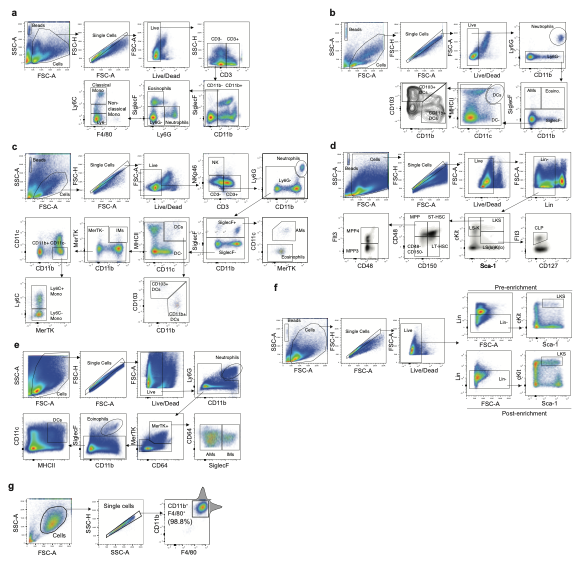

**Figure S1: Flow cytometry gating and sorting strategies employed for the characterization and purification of innate immune populations.** Panels show gating strategies for the identification innate cell subsets in **(a)** peripheral blood, **(b)** BALF, **(c)** lung and **(d)** bone marrow. **(e)** FACS-sorting strategy used for isolation of lung cell populations. **(f)** Magnetic enrichment of Lin^-^ populations from live bone marrow cells prior to downstream analysis. **(g)** Flow cytometry-based assessment of BMDM purity following *in vitro* differentiation.

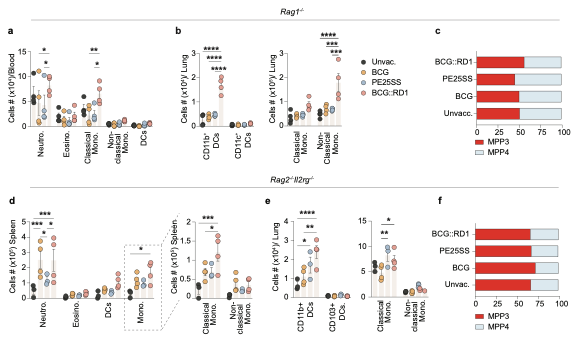
**Figure S2: Innate immune populations and bone-marrow multipotent progenitor composition in immunodeficient mice.** Panels **(a-c)** show data from *Rag1^-/-^* mice: **(a)** total innate immune cell counts in peripheral blood, **(b)** lung DC and monocyte subsets, and **(c)** % of bone-marrow multipotent progenitor subsets MPP3 versus MPP4. Panels **(d–f)** show data from *Rag2^-/-^Il2rg^-/-^* mice: **(d)** total innate immune cell counts in spleen, **(e)** lung DC and monocyte subsets, and **(f)** frequency of bone-marrow MPP3 versus MPP4. Statistical significance was calculated using **(a, b, d, e)** Two-way ANOVA with Tukey posthoc test (**p* < 0.05, ***p* < 0.01, ****p* < 0.001).

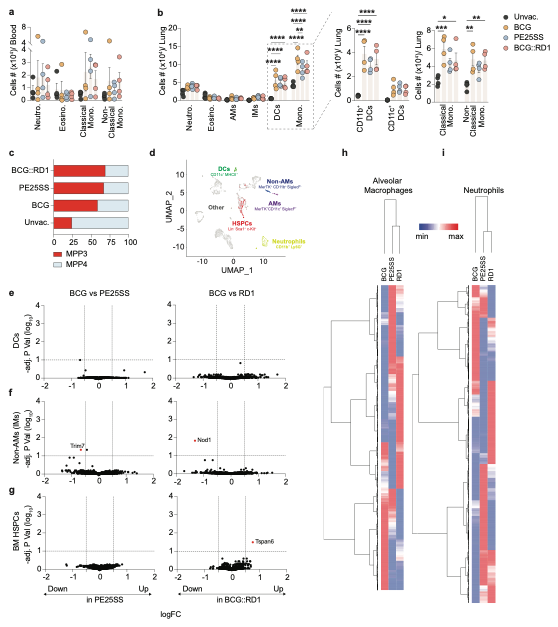

**Figure S3: Innate immune cell composition and single-cell transcriptional profiling following mucosal vaccination.** Total innate immune cell counts in **(a)** blood, **(b)** lung, and **(c)** % of bone marrow MPP3 vs MPP4 cells in C57BL/6 mice 30 days of p.v.. **(d)** UMAP visualisation of scRNA-seq data showing clustering of immune cell populations based on cannoical gene marker expression. DEGs idenified in **(e)** DCs, **(f)** IMs (non-AMs), and **(g)** BM HSPCs (LKS^+^). Heatmaps showing normalised expression levels for total genes expressed in **(h)** AMs and **(i)** neutrophils. Statistical significance was calculated using **(a, b, d, e)** Two-way ANOVA with Tukey posthoc test (**p* < 0.05, ***p* < 0.01, ****p* < 0.001).

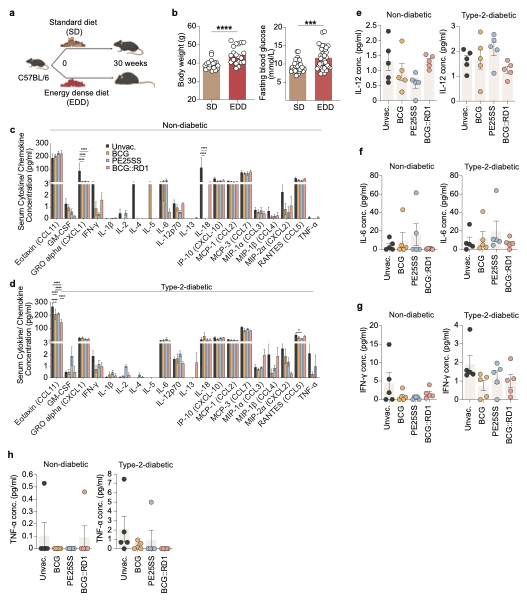

**Figure S4: Induction of diet-induced murine type 2 diabetes and systemic cytokine and chemokine responses following *Mtb* infection.** To induce T2D, **(a)** male C57BL/6 mice were fed an ED diet for 30 weeks and assessed for **(b)** body weight and fasting blood glucose following the diet intervention. Total serum cytokine and chemokine concentrations measured at 45 days p.i. in **(c)** non-diabetic and **(d)** type 2 diabetic mice. Comparison of key pro-inflammatory cytokines; **(e)** IL-12, **(f)** IL-6, **(g)** IFN-γ and **(h)** TNF-α. Statistical significance was calculated using**(b)** Student’s t-test, **(e-h)** One-way ANOVA with Tukey posthoc test or **(c, d)** Two-way ANOVA with Tukey posthoc test (**p* < 0.05, ***p* < 0.01, ****p* < 0.001).

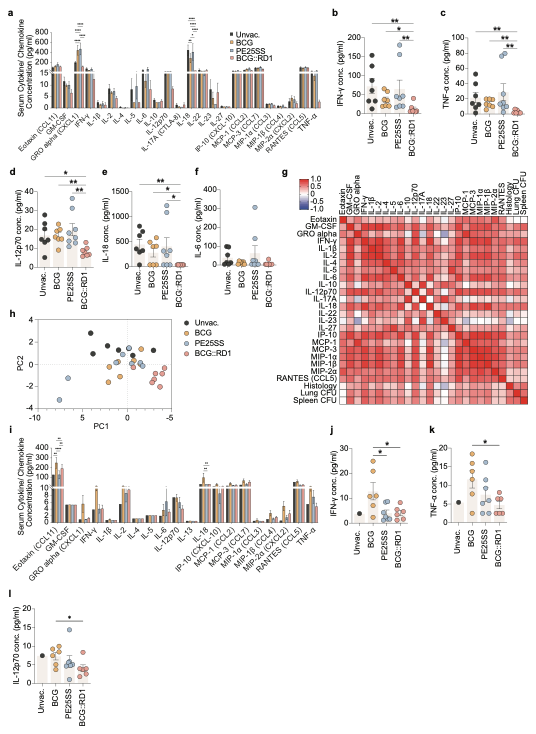

**Figure S5: Systemic cytokine and chemokine responses in immunodeficient mice following *Mtb* infection.** Total serum cytokine and chemokine levels measured at 45 days p.i. in **(a)** *Rag1^-/-^* and **(i)** *Rag2^-/-^Il2rg^-/-^* mice. Comparison of key pro-inflammatory cytokines; **(b)** IFN-γ, **(c)** TNF-α, **(d)** IL12, **(e)** IL18 and **(f)** IL-6 in *Rag1^-/-^* mice and **(j)** IFN-γ, **(k)** TNF-α, **(l)** IL12 in *Rag2^-/-^ Il2rg^-/-^* mice. **(g)** Correlation matrix and **(h)** PCA plot illustrating associations among serum cytokine levels, organ bacterial burdens (CFU) and lung tissue inflammation. Statistical significance was calculated using **(b-f, j-l)** One-way ANOVA with Tukey posthoc test or **(a, i)** Two-way ANOVA with Tukey posthoc test (**p* < 0.05, ***p* < 0.01, ****p* < 0.001).
